# Automated Deep Learning-based Multi-class Fluid Segmentation in Swept-Source Optical Coherence Tomography Images

**DOI:** 10.1101/2020.09.01.278259

**Authors:** Jonathan D. Oakley, Simrat K. Sodhi, Daniel B. Russakoff, Netan Choudhry

## Abstract

**Purpose:** To evaluate the performance of a deep learning-based, fully automated, multi-class, macular fluid segmentation algorithm relative to expert annotations in a heterogeneous population of confirmed wet age-related macular degeneration (wAMD) subjects.

**Methods:** Twenty-two swept-source optical coherence tomography (SS-OCT) volumes of the macula from 22 from different individuals with wAMD were manually annotated by two expert graders. These results were compared using cross-validation (CV) to automated segmentations using a deep learning-based algorithm encoding spatial information about retinal tissue as an additional input to the network. The algorithm detects and delineates fluid regions in the OCT data, differentiating between intra- and sub-retinal fluid (IRF, SRF), as well as fluid resulting from in serous pigment epithelial detachments (PED). Standard metrics for fluid detection and quantification were used to evaluate performance.

**Results:** The per slice receiver operating characteristic (ROC) area under the curves (AUCs) for each of these fluid types were 0.90, 0.94 and 0.94 for IRF, SRF and PED, respectively. Per volume results were 0.94 and 0.88 for IRF and PED (SRF being present in all cases). The correlation of fluid volume between the expert graders and the algorithm were 0.99 for IRF, 0.99 for SRF and 0.82 for PED.

**Conclusions:** Automated, deep learning-based segmentation is able to accurately detect and quantify different macular fluid types in SS-OCT data on par with expert graders.

## Introduction

Age-related macular degeneration (AMD) is a leading cause of blindness for people over the age of 50, a demographic that is an increasing percentage of the world’s population. Declining fertility rates and increasing longevity have been shifting the age distribution, in industrialized countries, towards older age groups.^1^ Approximately 170 million individuals are affected with AMD globally and prevalence of AMD is expected to increase to 288 million by the year 2040;^2,3,4,5^ by 2030 it is estimated that the number of cases in the US alone will reach 3.5 million. The prevalence of this and other ocular diseases places a tremendous burden on the healthcare system, and there is a critical need to streamline diagnosis and improve management of these patients.

Wet age-related macular degeneration (wAMD), the advanced, exudative form of AMD, can be managed using therapeutic agents, but relies on the assessment of fluid amount and type. Optical coherence tomography (OCT) revolutionized the management of AMD and has been used in investigative trials to link anatomical structure to function in the form of visual acuity (VA).^6^ This ability to measure thickness as a correlate of fluid amount, provides a quantitative value to measure treatment response. There have, however, been discrepancies between this value and reported functional outcomes in the literature.^7^ In the absence of automation, identifying the type and location of fluid is a qualitative clinical approach to determining disease activity and thus individualizing treatment.^8^ Manually assessing the OCT data in this subjective way relies on the clinician/grader’s skill and can be time consuming.

The field of computer vision offers a more efficient approach to analysis tasks involving automated segmentation algorithms to assess retinal thickness. These types of software packages are bundled with the OCT devices and remain instrument-specific. While they are able to accurately segment in normal patients, they tend to perform poorly in cases of pathology, particularly advanced disease states, where the need for this evaluation is paramount.^9,10^ Furthermore, retinal thickness measurements are only a surrogate measure of fluid volume, that is, fluid detection, quantification and classification are not presently in the armamentarium of analysis tools available clinically.

Deep learning has dramatically changed computer vision making complex tasks, such as automated fluid segmentation, feasible. One of the most powerful aspects of deep learning is that the framework automatically learns both the classifier as well as the salient features that feed into it. Prior to this innovation, feature extraction typically required meticulous manual intervention. We apply this approach in this study using convolutional neural networks (CNNs) and supervised learning. The CNNs are capable of building rich, layered (deep) representations of the data which are then classified through additional layers of representation and learned associations. Such significant technical improvements have advanced to levels matching expert human graders, taking seconds to perform tasks that would otherwise take hours. This functionality is of significant interest, particularly in ophthalmology where powerful imaging devices are relatively inexpensive, the device software limited, and image data is key to clinical management. However, ahead of clinical deployment these algorithms will require extensive evaluation and, ultimately, regulatory approval.

In the following we report on an initial study of the performance of macular fluid segmentation in a heterogeneous, wAMD population. Previously reported studies include Schlegel et al., who report on fluid segmentation in AMD, diabetic macular edema (DME) and retinal vein occlusion (RVO) cases, using two different OCT devices.^11^ Mehta et al., similarly report on an approach for intra-retinal fluid (IRF) segmentation that showed an existing deep learning model’s improvement relative to manual graders after transfer learning.^12^ More recently, and underlining the interest of this particular application, an on-line challenge was organized stemming from a medical imaging conference that assimilated and compared a number of different deep learning approaches for segmenting three retinal fluid types in OCT data; intra- and sub-retinal fluid (SRF) and pigment epithelial detachment (PED).^13^ This task maps nicely to the work we present in the following and is returned to in the discussion section of this paper.

## Subjects and Methods

This study was Institutional Review Board approved and followed the tenets of the Declaration of Helsinki. Written informed consent was obtained from all participants and was in accordance with current ICH/GCP guidelines, section 4.8.10.

The data was collected using the Topcon Triton SS-OCT device (Tokyo, Japan), using the macular protocol, the scan was centered on the fovea and covered a lateral area of 7mm^2^, with a depth of 2.58mm. The data used in this study was acquired from 22 patients where each volume comprised 256 B-scans, with each B-scan being 512-by-992 pixels. The key inclusion criteria were: (1) patients had to be ≥ 25 years old; (2) diagnosed with wAMD and (3) wAMD treatment naíve.

### Manual Segmentations

Two expert graders, SS and JO, separately annotated every fourth B-scan using custom image labeling software. Consensus segmentation results were arrived at in final consultation and adjudication with a senior retina specialist (NC). This created volumes with every 2nd B-scan manually labelled. Adopting a similar sparse approach as input to generating additional labeling data from De Fauw et al., the intermediate B-scans were segmented using the same semantic segmentation algorithm reported on for the final result, where the training data was every 2nd B-scan from the same volume (Figure 1)^14^. The final correction of the automated labeling was done manually, ultimately ensuring that all 22 volumes (5,632 B-scans) used in the study were fully labeled and could be used as ground truth segmentations.

**Figure 1.**
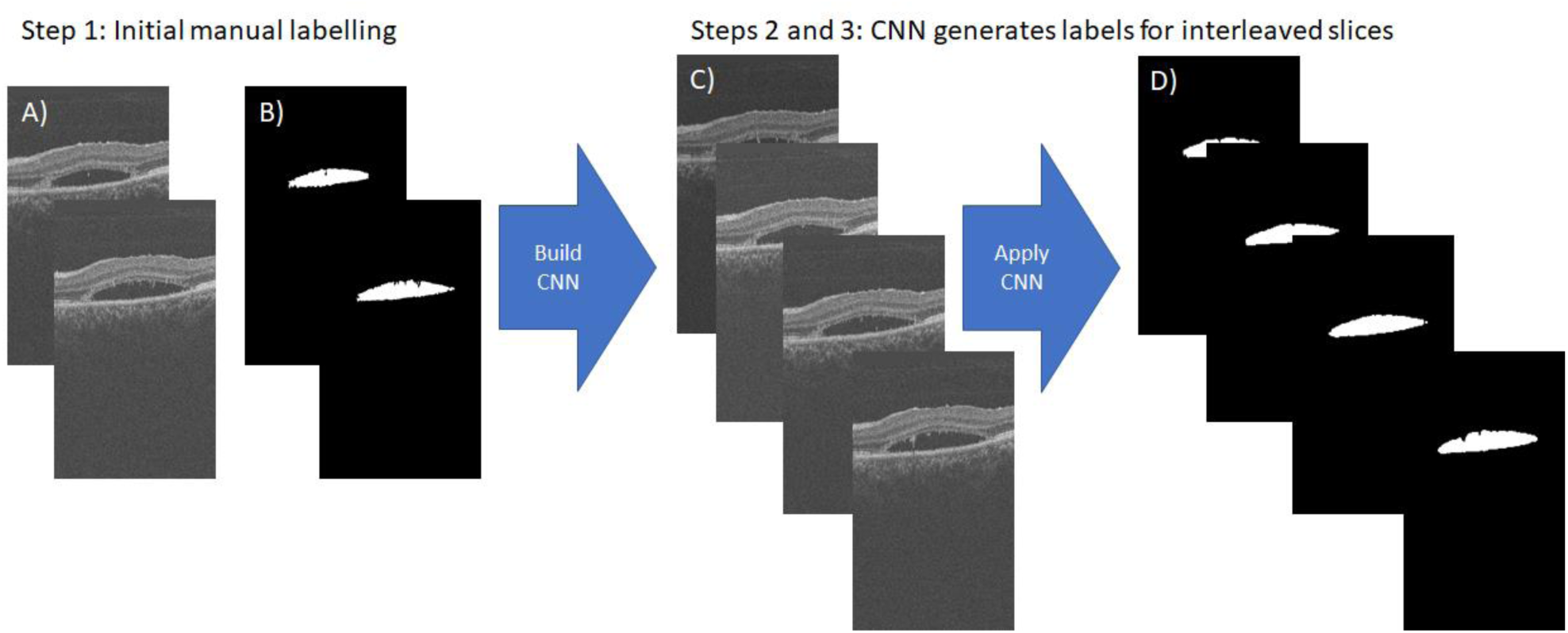
Use of a CNN to generate additionally labeled images that were then subsequently verified manually. For manually labeled images in the training set (Step 1), a CNN is capable of accurately labeling the neighboring images when presented in the test set (Steps 2 and 4). Example input images (A) are used with associated manual labels (B) to create an initial CNN. Unlabeled images (C) from the same volume are used as test data applied to the CNN. The output labels (D) are then manually verified such that they may themselves be used as ground truth.

### Pre-Processing

Each input volume’s retinal layers were automatically segmented by Orion™ version 3.0 (Voxeleron, San Francisco, USA), which is research software that performs OCT analysis. This segmentation information was then encoded into a separate channel that was later entered into the supervised learning process. Existing literature has used, separately, layer information as features in support of the classification of fluid pockets,^15^ in their words leveraging the fact that retinal fluid is known to exhibit layer dependent properties and thus is used by the model to help determine the type of fluid; IRF typically being higher up in the volume, SRF close to and above the RPE, PEDs below the RPE, but not below the retina itself. In Figure 2, the column of images on the left represent single B-scans within the volume, that on the right are the layer segmentation result encoded as image data for each of those B-scans. Lastly, each B-scan and the associated label images were sub-sampled to 256-by-256 pixels in size.

**Figure 2.**
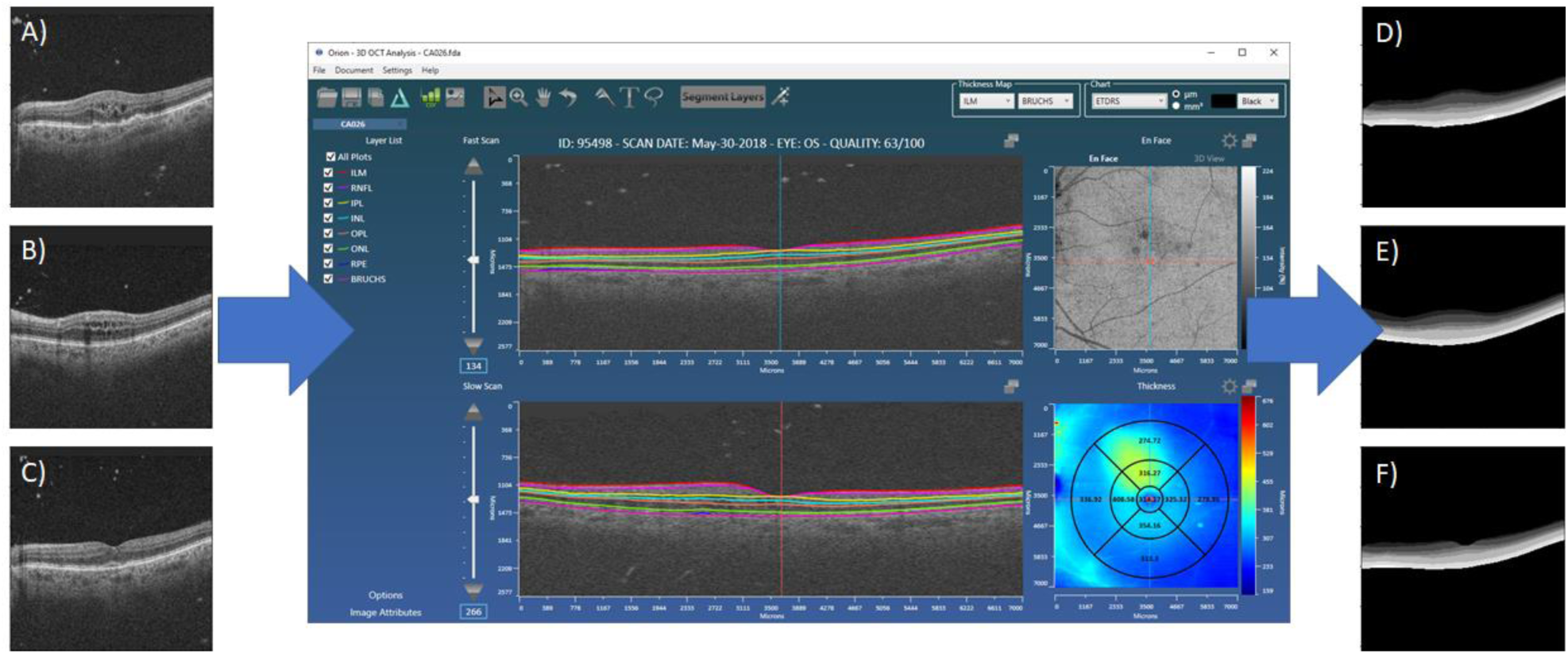
Each image volume is first automatically segmented using Orion into 8 retinal interfaces. This results in 7 layers that can be encoded into image form and used as an additional channel in model creation, thus encoding spatial information regarding the location of fluid within the retina. Example B-scans are shown in (A), (B) and (C). They are taken from the volume shown loaded into the Orion software application (center). The resulting segmentations are shown in (D), (E) and (F), respectively.

## Deep Learning

### Method

Given 22 volumes, the study comprised a total of 5,632 B-scans, and due to pre-processing, each 2D B-scan was 256-by-256 pixels. For the task of supervised learning, a cross validation (CV) approach was adopted for training and subsequent testing, where data was very strictly stratified by subject. Given the limited number of patients, we used 10-fold CV, so on each fold a model was built for all but two or three subject eyes. During the learning process, the model was assessed based on a validation set that randomly chose 5 subjects from the training data (1,280 B-scans). Therefore, on average, training data comprised data from 15 subjects and validation data came from the remaining 5 subjects. This strict stratification to subject eye means that, while the learning uses only training set, we simultaneously validate the model on truly unseen data, stopping the learning process only when the validation error is clearly increasing. This is a standard technique in supervised learning that is used to prevent over-fitting. It does, however, require that the optimization and learning rate are chosen appropriately.

The training data was augmented using random scaling, rotations and shifting such that, for every input B-scan, six additional B-scans were generated. We did not augment the validation data.

### Implementation

The deep learning architecture used for first the training and then the testing was that of the U-Net, which is a version of the autoencoder that uses skip connections to better maintain detail across different scales^16,17^. The implementation uses Tensorflow 2.0 (Google, Mountain View) and training and testing were performed on a Dell Precision 7920 workstation, running NVIDIA Quadro GV100 GPUs. The network is fed the input 2D images and their encoded segmentations in a second channel. At the same scale, the labels are applied as input (see Figure 3). The network uses three encoding levels, as shown in Figure 4, and three decoding levels. Learning is based on minimizing the model’s loss, where the loss function uses categorical cross entropy (CCE) with an added term that weights a different measure called the Dice similarity coefficient (DSC). The DSC, for two sets is twice the area of their overlap divided by the total number of elements in both sets. CCE is used for multi-class classification where the labels use one-hot encoding and defines a measure of dissimilarity between the true value and that which is predicted. In our application, non-fluid, or background regions will dominate this cost function that is to be minimized, requiring appropriate weighting of the fluid labels to be applied. This was achieved through the additional weighted term to our cost function that maximizes the total dice coefficient score across the labels. The weighting can then be set so as to diminish influence of the background pixels on the overall dice score. Empirically we look for loss values to decrease with increasing epochs in a fairly monotonic way; i.e., to ensure that, on the unseen validation data, the network continued learning without over-fitting. This combination of CCE and weighted dice loss works well and enables a fairly continuous reduction in validation loss with epochs. There are, however, other hyper-parameters are at play: drop-out was set to 0.25, batch normalization was applied, and the batch size was 8. The optimizer used was Adadelta^18^.

**Figure 3.**
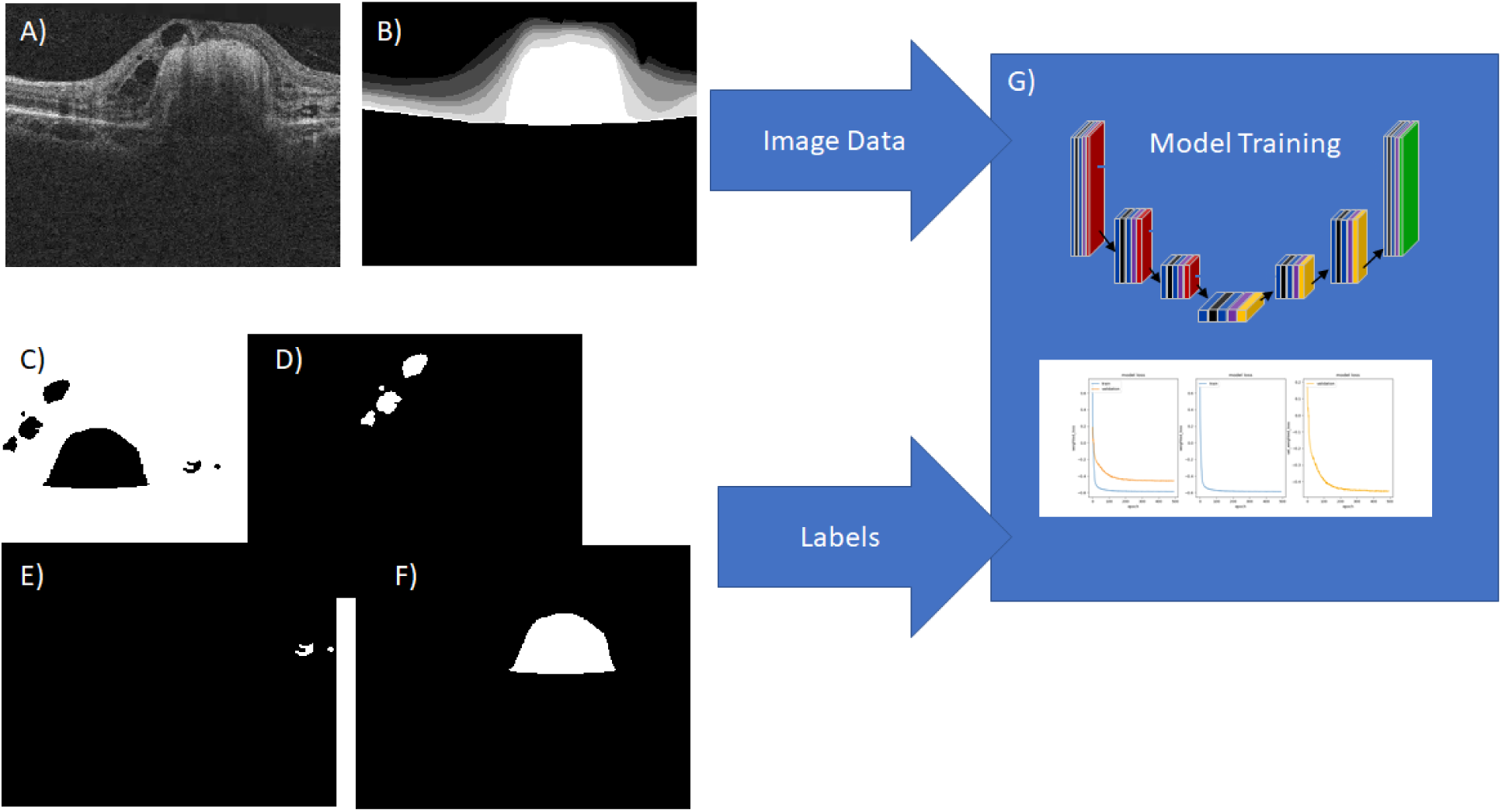
The supervised learning approach using a deep convolutional neural network architecture. The input image data are the B-scans (A) and their associated encoded segmentation results (B). The input labels are one-hot encoded versions of each fluid type. This example has one of each: background (C), IRF (D), SRF (E) and PED (F). Typically, one or more of the fluid label images will be entirely blank, meaning that fluid type was not present in that B-scan. G) portrays the deep learning architecture that takes in the data and labels and minimizes the loss between the two for the training data. An example plot of loss value decreasing over iterations is shown, where the training loss is shown in blue and the validation loss in orange.

**Figure 4.**
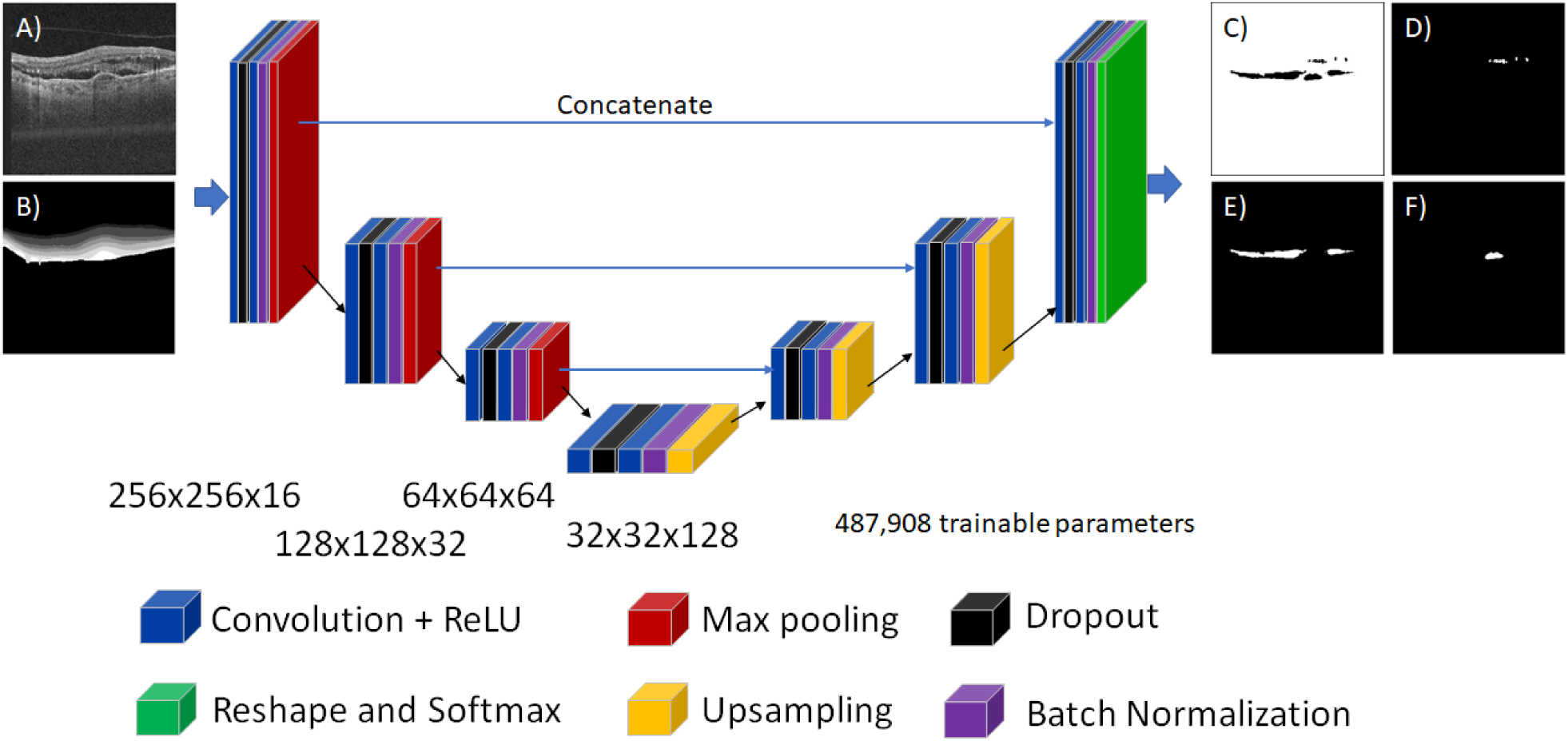
The network architecture used in these analyses. For the two-channel input image (A,B), it outputs four images giving the probability score, at each pixel, of belonging to one of the four classes: background (C), IRF (D), SRF (E) and PED (F). Dropout was set to 0.25 and the final softmax layer ensures these are normalized and can thus be interpreted as probabilities.

The DSC for a single label is defined as:

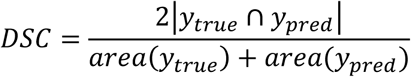

where *y*_*true*_ and *y*_*pred*_ are the labels and the current prediction of the labels, respectively. The weighted dice term, DW, then becomes:

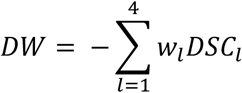

where *l* denotes the labels – background, PED, SRF and IRF – and *w*_*l*_ denotes the weights applied to each of these. The final loss function then becomes:

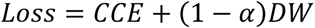

where α controls the influence of the weighted dice term and is set to 0.2. The weights used are *w* = [0.05, 0.5, 0.225, 0.225], which was set *a priori* based on an estimate of relative proportions in wAMD. It should be noted that if the first weight term is too high, background will tend to dominate the segmentation. The distribution of the other weights is less critical.

The optimizer’s initial learning rate (ILR) was set to 0.5, which changed every *n*^*th*^ iteration such that, at epoch *i* the learning rate used is given as:

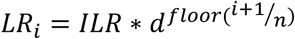

With *d* set to 0.95, the drop fraction, this resulted in an exponential decay of the initial learning rate that stabilized to a minimum near 600 epochs. Training stops when the loss score applied to the validation data set stops diminishing for 50 epochs.

## Results

### Fluid Detection

Fluid detection is reported on a per B-scan and per volume basis using receiver operating characteristic (ROC) curves, with the summary statistic being the area under the curve (AUC). For a given B-scan, the network outputs a probability score for each class, facilitating the plot of ROCs and measure performance. In evaluating on a per volume basis, we were are only able to report IRF and PED performance as all 22 wAMD subjects had at least some SRF as reported by the manual labelling, meaning no negative examples were available. These are given in Figure 5, with precision and recall values in Table 1.

**Table 1.** Precision and recall proportions for each fluid group by B-scan and volume. Precision measures the proportion of positively labeled cases that were actually correct, and recall measures the proportion of positive cases that were identified correctly. Precision example: when the algorithm says a volume contains PED, it is correct 97.8% of the time. Recall example: our algorithm correctly identifies 92.1% of volumes with SRF.

|  | Per B-scan |  |  | Per volume |  |  |
| --- | --- | --- | --- | --- | --- | --- |
|  | IRF | SRF | PED | IRF | SRF | PED |
| **Precision** | 0.503 | 0.749 | 0.729 | 1 | 1 | 0.978 |
| **Recall** | 0.769 | 0.827 | 0.842 | 0.823 | 0.921 | 0.898 |

**Figure 5.**
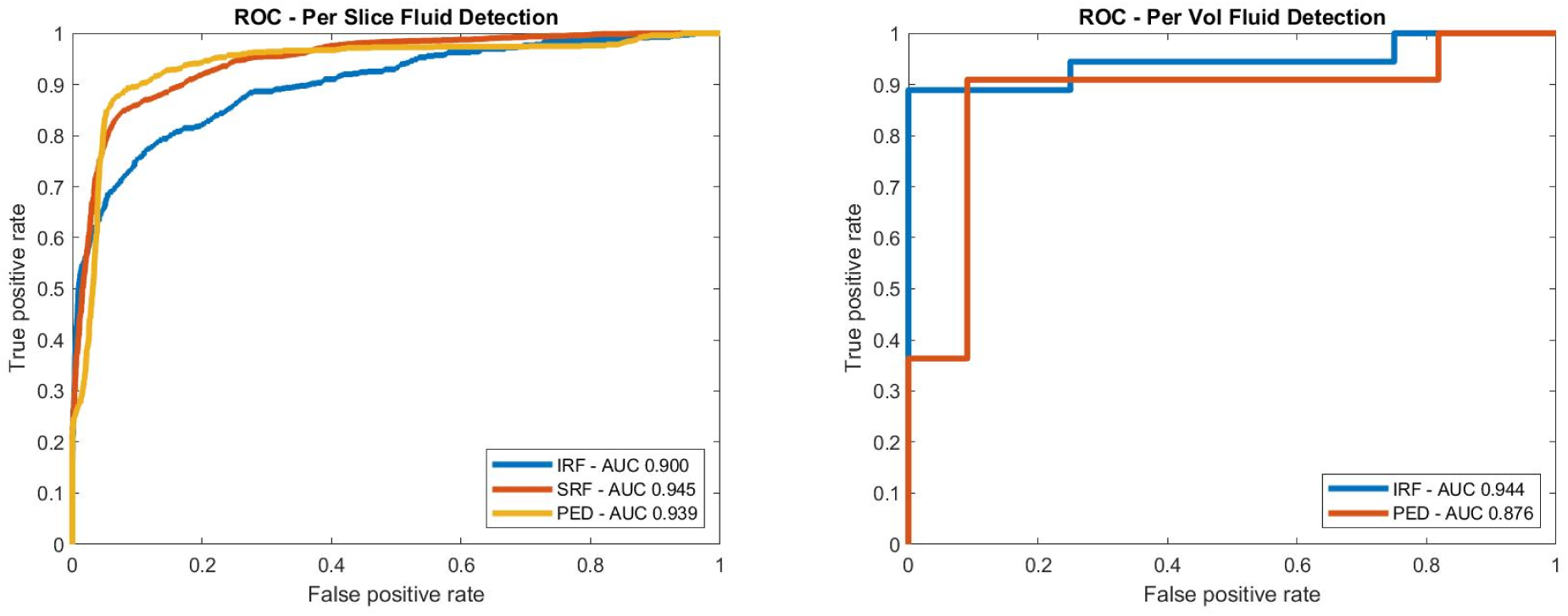
Per B-scan AUCs were 0.900, 0.945 and 0.939 for IRF, SRF and PED, respectively. Per volume results were 0.944 and 0.876 for IRF and PED.

Precision is defined as the number of true positives (TP) divided by the total number of *reported* positive cases, true and false (FP): TP/(FP+TP). Recall is TP divided by the total number of *actual* positive cases: TP/(TP+FN). The values are calculated at the Youden index of the ROC curve.

### Fluid Quantification

Given manual segmentations of all B-scans in all volumes, we report the ability to quantify fluid by directly correlating the total volume of fluid for each of the 22 subjects that was manually delineated with that automatically determined. The coefficient of determination (R^2^) is used as a summary statistic for correlation, and both the correlation and Bland-Altman plots are shown in Figure 6. Furthermore, the area of overlapped, as quantified by the DSC, between graders and the algorithm was examined (Figure 7). Finally, example result images are given in Figure 8 and Figure 9.

**Figure 6.**
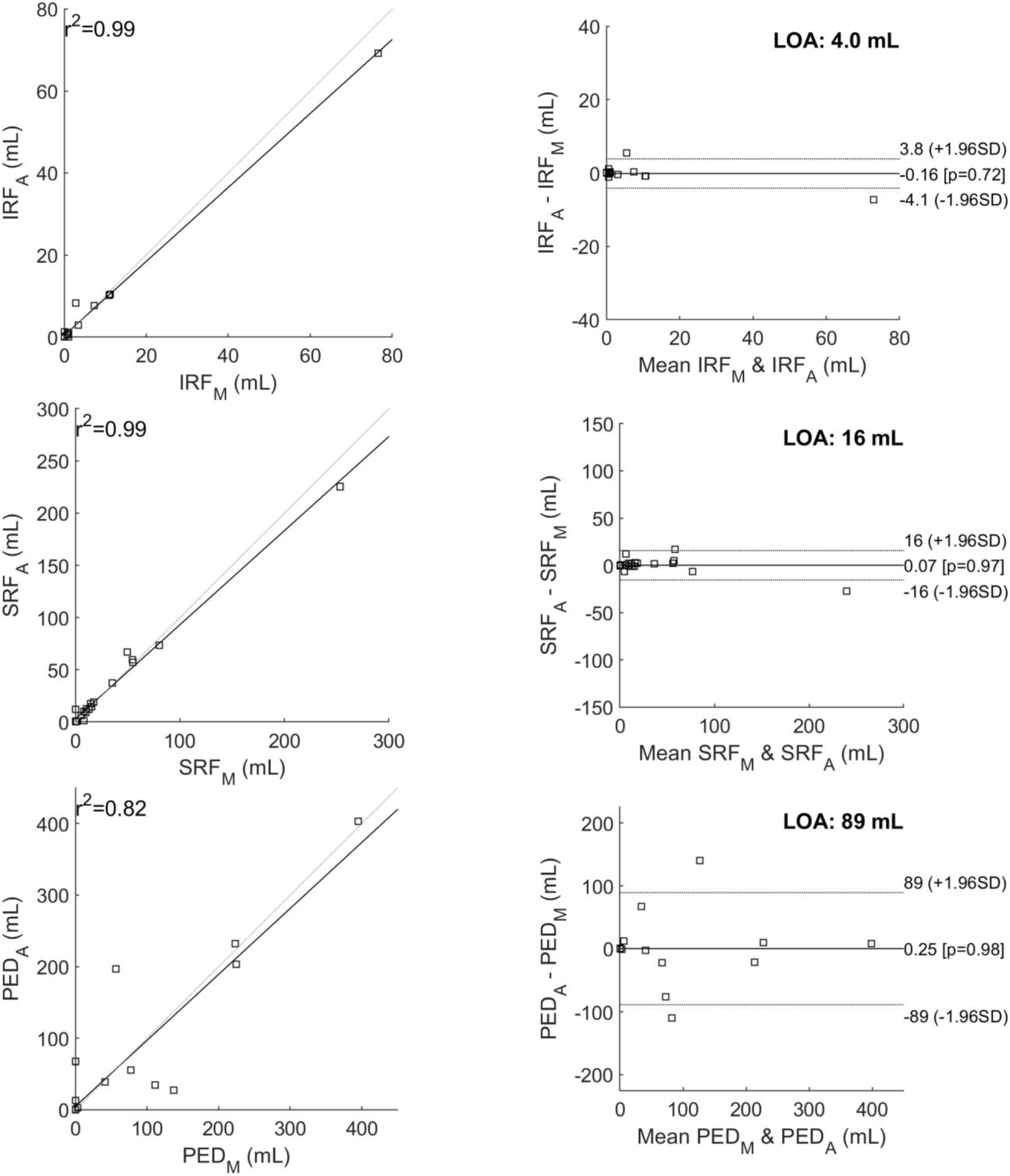
The above shows the manual versus automated reported total areas for each fluid type across all 22 volumes (left) and the Bland-Altman plots on the right. The correlation scores are 0.99, 0.99 and 0.82 for IRF, SRF and PED, respectively (top to bottom). For the Bland-Altman plots, the manual values are denoted with ‘_M’ and the automate values with ‘_A’. Narrow limits of agreement are shown for IRF and SRF, but are larger for PED, as we address in the discussion.

**Figure 7.**
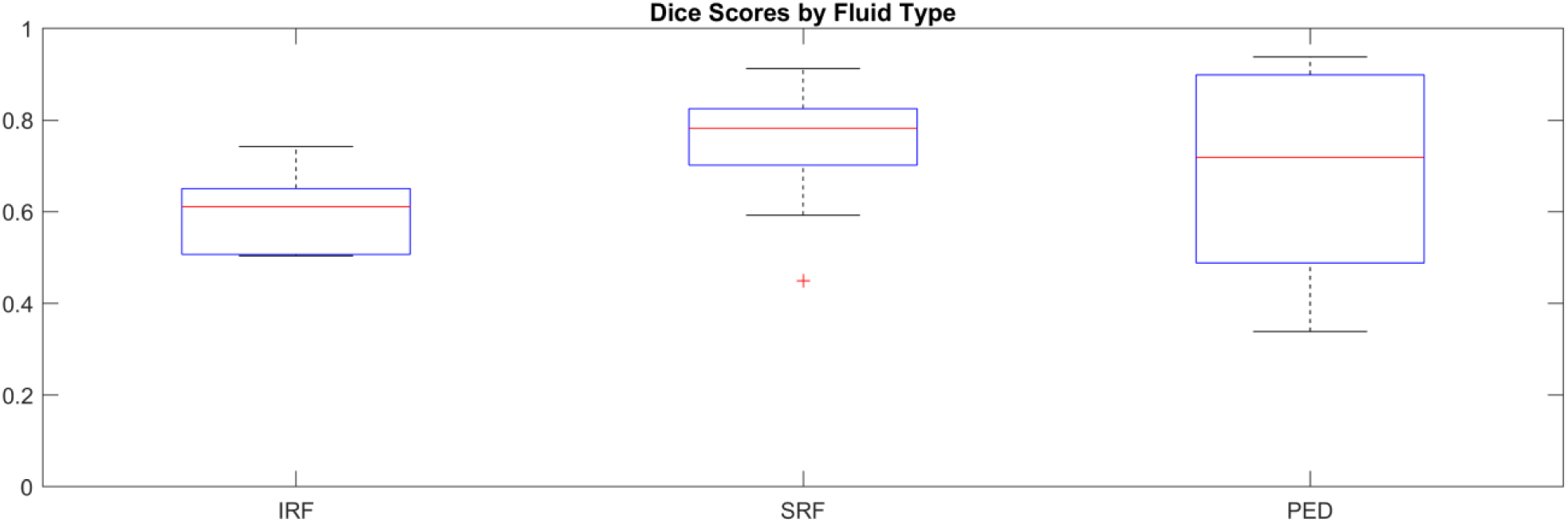
Dice coefficients for all cases where a given fluid type was manually labeled as being above 1mL.

**Figure 8.**
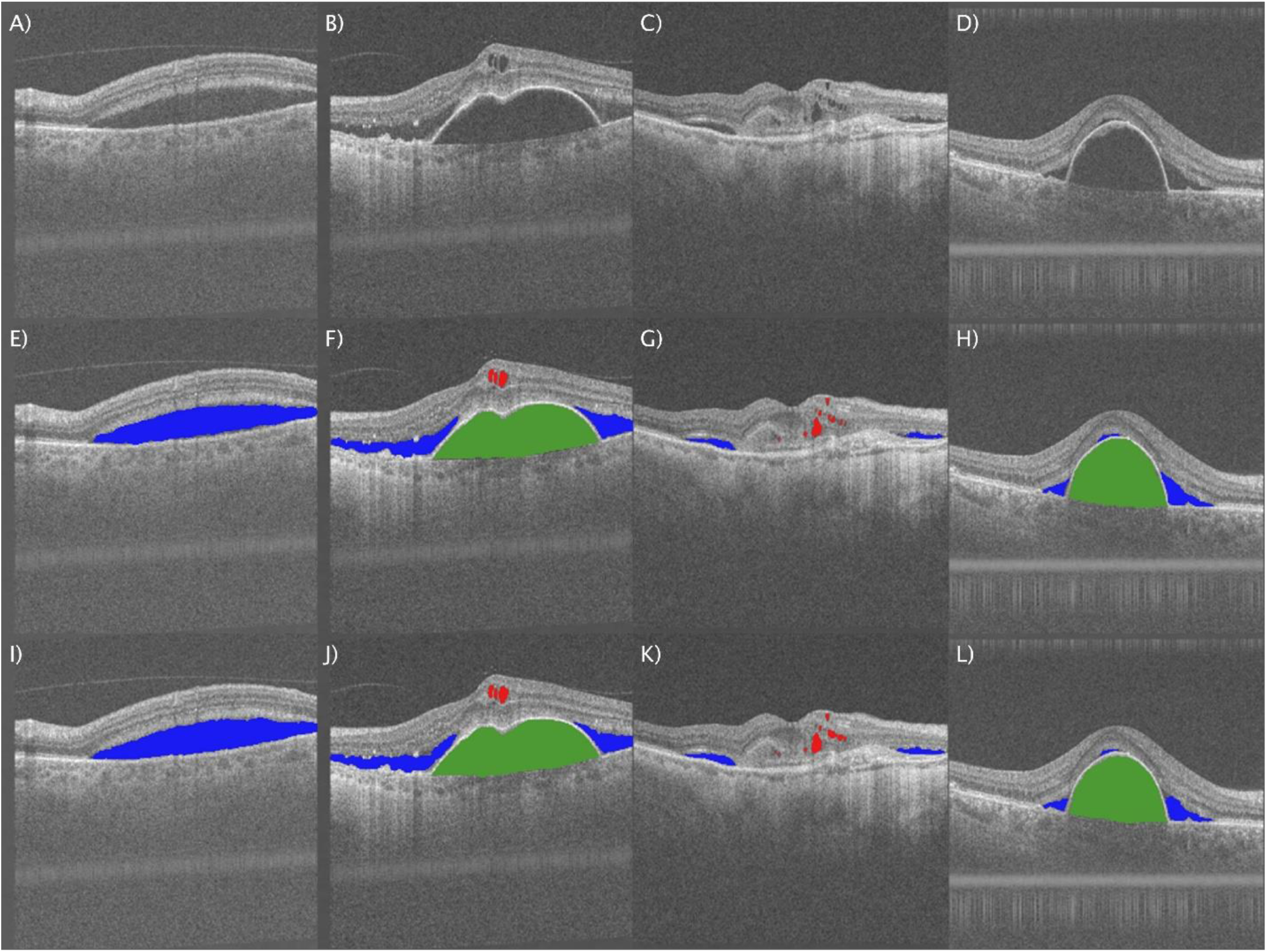
Example segmentations. Input B-scans (A,B,C,D), manual segmentations (E,F,G,H), and the automated segmentations (I,J,K,L). IRF is labeled red, SRF in blue and green constitutes PED.

**Figure 9.**
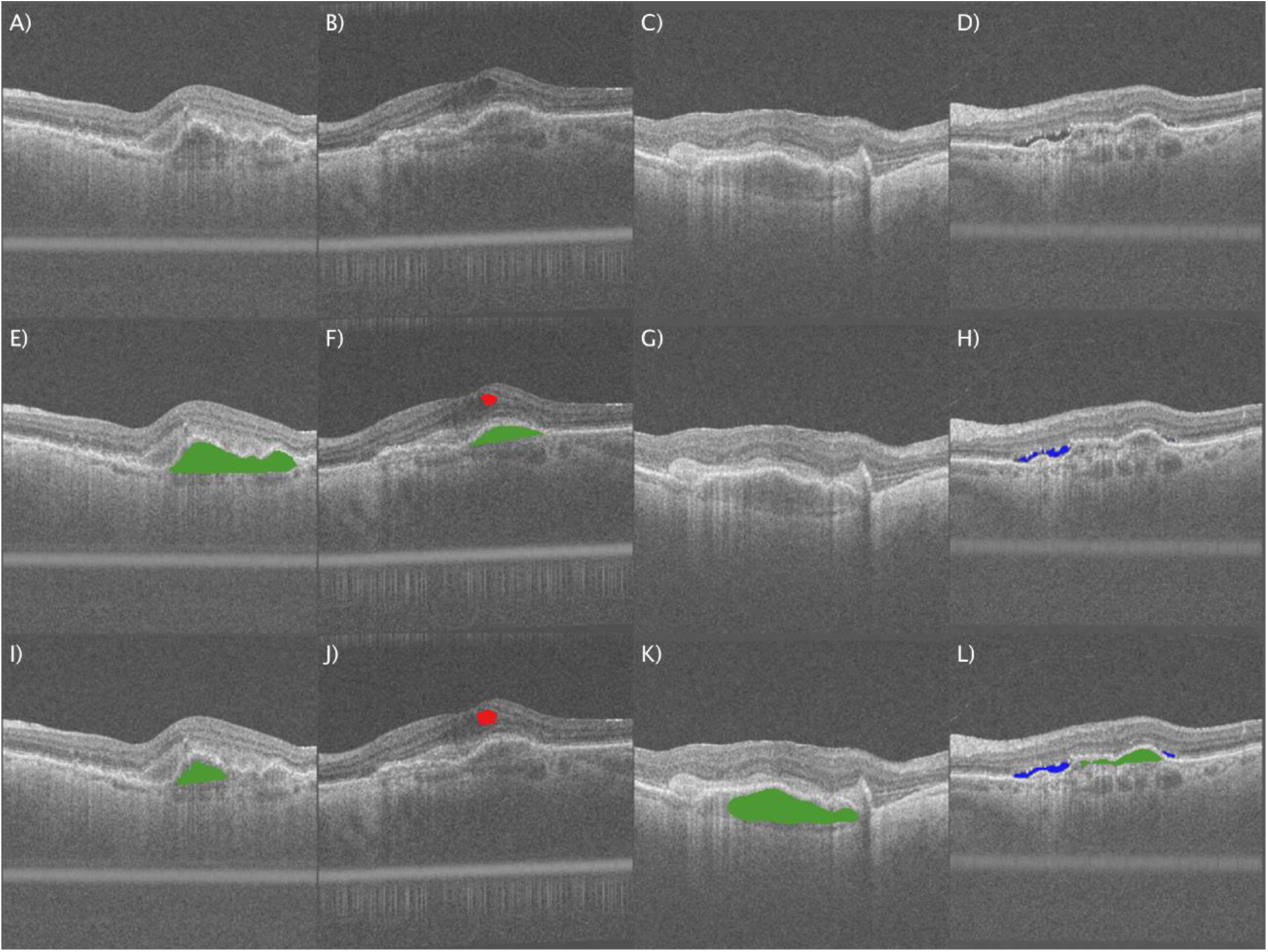
The above shows example input B-scans exhibiting fibrovascular PEDs (A,B,C,D). The manual labelings below (E,F,G,H) decided the only (E) and (F) contained fluid and (G) and (H) were mostly fibrous tissue. The automated segmentations show that cases (I), (K) and (L) labeled as PEDs, but not (F).

## Discussion

We have shown that the performance of automated segmentation of three fluid types in SS-OCT is in very strong agreement with that done manually by expert graders. To the best of our knowledge, this is the first study to report on such a technique using SS-OCT. Previous work using deep learning has, for example, reported on the segmentation of the retina and choroid and also disease detection in SS-OCT, but not retinal fluid segmentation.^19,20^ As an imaging modality, SS-OCT is becoming clinically more relevant than the more conventional and established spectral domain OCT (SD-OCT) due to a number of distinct advantages. A first is the speed of acquisition that is typically four times as fast as SD-OCT devices – the Triton device used here acquires 100,000 A-scans per second – which allows for more data to be captured over wider fields of view in a short amount of time. Prototype devices now operate at up to 400,000 A-scans a second.^21^ Speed of acquisition minimizes patient chin rest time and reduces the amount of motion in the scan. A second advantage is the longer wavelength of the light source. This allows the light to penetrate deeper into the retina, supportive of better visualization of the choroid and structure below the RPE. And while SS-OCT has slightly reduced axial resolution relative to similar SD-OCT devices, the overall signal to noise ratios are higher.

Deep learning, however, is the enabling technology of this study, prior to which earlier approaches are quite varied. One good example from Bogunović et al. assimilates a number of proven techniques to achieve fluid segmentation.^15^ Their approach uses layer segmentation to constrain the segmentation region and also support positional information about the fluid that is used, along with textural features, to assign probabilities of voxels within the top and bottom segmented layer, as “fluid” or “no fluid”. The probability assignment uses an artificial neural network (ANN) as the classifier where supervised learning is based on a leave-one out (LOO) CV approach for the 30 volumes used. This soft segmentation is hardened in a final step that uses fully 3D geodesic graph cuts to isolate each of the fluid pockets producing binary labels.

Similar methods exist where layer segmentation is initially used to constrain the problem and then a number of techniques such as region growing, or a combination of thresholding and boundary tracing or clustering based segmentation to arrive at the final result.^22,23,24,25^

An important early deep learning reference is the work of Roy et al. who compare a variety of U-Net implementations as well as different loss functions on 110 labeled images from 10 subjects.^26^ Careful to stratify by patient, subjects 1 through 5 were used for training, 6 through 10 for testing, with further evaluation using 8-fold CV. Other previously reported studies include Schlegel et al. who report on fluid segmentation in AMD, diabetic macular edema (DME) and retinal vein occlusion (RVO) cases.^11^ This was performed using two different OCT devices and has become an important reference. For comparison, considering the AMD cases only, fluid types (IRF and SRF) were segmented using a deep CNN and both 10- and 4-fold CV. The number of volumes were 70 and 257, respectively for the two different instruments. In general, the larger the data set, the smaller the number of folds that may be used in assessing performance. Performance is impressive, with their volume-based detection of fluid for AMD had AUCs of 0.95 and 0.98 for IRF and SRF with the Cirrus device (Carl Zeiss Meditec, Dublin, USA); and 0.93 and 0.90 with the Spectralis device (Heidelberg Engineering, Heidelberg, Germany). Mehta et al. similarly report on a deep learning approach for here one kind of IRF segmentation.^12^ Their reporting mechanism was to show that the DSC between the automated algorithm and the manual graders improved after transfer learning. This is expected given that the original model was trained on data from a different instrument: the model used, from Lee at al. was created based on Spectralis data and was then further trained for Cirrus data using 400 OCT B-scans (2D images) from 58 patients.^27^ It was deployed on a test set comprising of 70 scans from 70 different patients. The architecture was necessarily that developed by the original authors as transfer learning was applied. Results showed that the distribution of DSCs improved after transfer learning as a result of their approach.

The work of Schlegel is of particular importance given its recent deployment in a retrospective clinical analysis that was able to correlate fluid amounts to clinical outcomes.^28^ A key difference in our study, however, is the additional segmentation of PED segmentation within the same network. This becomes feasible with this data set, because of the method of spatial injection that uses the layer segmentation software (Orion™). This we see is also the case in the analysis of three fluid labels in the RETOUCH challenge where, in the best performing instances, the approach is to first segment the data, second train a CNN with image data and segmentation as input, and third refine the final result using an additional classifier.^13,29^ We see a benefit of having the segmentation element using non-trained data and all of the fluid segmentation within a single CNN, as is reported here. But consensus exists that encoding spatial information about the retinal tissue as an additional channel to the input image data is shown to improve the overall performance of the segmentation algorithm. While fluid segmentation alone can be accomplished using just the input image data, the additional classification of each fluid type benefits significantly from this technique.

This pilot study is not without its limitations. Firstly, we have a limited number of subjects, meaning that, in CV analysis, there are only limited examples of a given fluid type in the training data when testing against a given subject. This is further confounded by our approach of splitting subject IDs across training, validating, and testing sets in an effort to ensure the network is learning but in a way that will generalize to unseen cases. This translates in modern parlance to ensuring that the variance is not so high and the model is not just overfitting the training data. The numbers are similar, however, to Bogunović et al., but as they have three volumes per subject, their leave-one out CV analysis might be unduly influenced by having subject data in both the train and test sets.^15^

While the lack of data may be a limitation of the analysis, it is not necessarily a limitation of the method. The deep learning paradigm thrives on data; the more data we have, the larger the learning space covered, meaning it’s more likely we have relevant examples to unseen data and the more natural the regularization of the resulting network supporting lower variance and a strong ability to generalize. In this respect, we look to the work of Schlegl et al. where the subject data is far larger.^11^ In considering just their population of wet-AMD patients, however, we see similar ratios of train/test splits to them, but they are able to apply this to scale. We consider this reference as a current standard, and as such note that our results are comparable.

Whenever supervised learning is applied, one must ensure that the ground-truth is correct. This can be challenging as the manual quantification of fluid presence, amount and type is difficult, extremely time consuming and highly variable across graders. As illustrated in Figure 9, we found a consistent definition of sub-retinal pigment epithelial (RPE) fluid in serous PED difficult, and this is a limitation as this can additionally lead to questioning the ground truth data used in 1) the training of the models using supervised learning and 2) the validation based again on manual labeling. Repeatability between manual graders has been previously reported as low, and in the context of segmentation tasks this is highlighted in the work of Bogunović et al. where their inter-observer analysis showed poor overlap between graders resulting in their ground truth being modified to be the union of the two graders.^30,15^ We have attempted to minimize the variability across graders by not reading blinded to each other, instead labeling intermittent slices of the same volume. This allowed each grader to read the images in the context of the other grader’s labelling. PED cases, however, can manifest as either serous, fibrovascular or drusenoid; or indeed a mix of these. Our target labeling was serous, which is sharply demarcated given the large fluid, dark (hyporeflective), regions and dome-like in shape. Fibrovascular PEDs on the other hand contain fibrovascular tissue, which refracts the OCT signal and shows, therefore, as lighter than fluid. Drusenoid PED are smaller, in general, creating an undulating RPE surface, and contain hyaline protein deposits, denser tissue that results again in a brighter appearance than fluid. Consistent labeling of our target detachment type, serous, was important, but it is still challenging to find a consistent manual “threshold” when some fluid only cases show clear signs of fibrovascular tissue; or vice versa. This is particularly evident in Figure 9 where fluid has been manually labeled in (E) and (F), but not in (G) and (H). Consequently, the algorithm too has difficulty in finding this difference. Ideally, one might iterate to find a consensus labeling in conjunction with the algorithm results, but that is also time consuming and such a consensus would ultimately improve but could also inappropriately bias the results. With enough data, however, the differentiation of PED subtypes is undoubtedly possible as is indicated in previous work.^14^

In conclusion, our study demonstrates that automated fluid detection and segmentation correlates highly with manual grading performed by experts. Combined segmentations of IRF and SRF with sub-RPE fluid in serous PEDs, offer a near complete automated solution for retinal fluid in wAMD. An automated approach to multi-layer fluid segmentation offers an advantage to physicians in a amidst a busy clinical setting and improved accuracy in managing patient care, particularly with the growing disease burden of wAMD. Furthermore, applying this deep-learning approach in clinical trials for therapeutic drug development could accelerate trial end points and the achievement of novel therapeutics within commercial timelines.

